# Unravelling the Oncogenic Potential and Prognostic Significance of *CKS1B* in Human Lung Adenocarcinoma and Squamous Cell Carcinoma: A Comprehensive Computational Analysis

**DOI:** 10.1101/2023.04.28.538730

**Authors:** Abu Tayab Moin, Md. Asad Ullah, Nairita Ahsan Faruqui, Yusha Araf, Tanjim Ishraq Rahaman, Kajol Ahsan, Md. Omer Faruq, Robiul Hasan Bhuiyan, Mohammad Jakir Hosen

## Abstract

Lung cancer (LC) confers to radical malignancy with a limited recourse of therapy worldwide. Consequently, LC has become the leading cause of cancer deaths in both men and women globally. Non-small cell lung cancer (NSCLC), one of the major LC types and accountable for a greater share of these cancer-associated deaths, further branches out to adenocarcinoma (LUAD) and squamous cell carcinoma (LUSC). A dearth of evident clinical symptoms coupled with the diagnosis feasibility only after advanced metastasis raises the need for precision in treatment apart from the existing chemical drug treatments. Precise guidance can be entailed by targeted therapies, utilizing the potential and thoroughly evaluated differentially expressed genes of cancer under speculation for tumor treatments. Cyclin-dependent kinase regulatory subunit 1B (CKS1B), a member of the conserved cyclin kinase subunit 1 (CKS1) protein family, regulates the cell cycle. Increasing evidence revealed that up-regulation of the *CKS1B* gene is associated with multiple human-related cancers, indicates its potential use as a targeted therapeutic for early detection and treatment. *CKS1B* has been found to be associated with poor prognosis in both LUAD and LUSC, while the prognostic significance of CKS1B in other types of cancer is not well established. Herein, we have performed a comprehensive bioinformatics analysis of factors involved in LUAD and LUSC with *CKS1B* and discussed its role as a potential biomarker for early lung cancer detection and treatment. While these evaluations demonstrate the immunotherapeutic features and prognostic value of *CKS1B*, further *in vivo* and *in vitro* studies are required to determine the accuracy of final applications.

## 1. Introduction

Cancer constitutes an array of diseases that entails the continual unregulated proliferation of abnormal cells, followed by its dissemination to surrounding tissues (1). The overall burden of cancer incidence and mortality has rapidly grown worldwide leading to an estimated 19.3 million new cancer cases and almost 10 million cancer deaths in 2020 (2). Cancer is the second leading cause of global deaths, responsible for about 1 in every 6 deaths. From the total number of cases, lung cancer (LC) accounts for about 11.6%. Non-small cell lung cancer (NSCLC) and small cell lung cancer (SCLC) are the two types of LCs among which NSCLC is the predominant form, making up about 80-85% of the total LCs (3). In contrast, the two major histologic subtypes of NSCLC include lung adenocarcinoma (LUAD) and lung squamous cell carcinoma (LUSC), comprising 40% and 30% of new lung cancer cases every year, respectively (4). At an early age, NSCLC patients usually exhibit no obvious clinical symptoms and with an additional lack of biomarker sensitivity and effective tools for prior detection, more than 75% of the NSCLC patients are still diagnosed at advanced stages with distant metastases (5). Moreover, the 5-year survival rate of LC was 17.8%, which is significantly lower than other cancer types (6). Despite improvements in novel therapies for LC patient survival, the 5-year survival rate was still less than 15% for advanced NSCLCs and was up to 80% for the initial stage of NSCLCs (7). Thus, an increased survival rate and reduced mortality rate can be attained through the early diagnosis and screening of lung cancer patients.

An up-regulation of differentially expressed genes (DEGs) could potentially act as biomarkers of LCs like LUAD or LUSC (8). An uncontrolled tumor cell growth, genome change, healthy cell damage as a result of the invasion of tumors into nearby tissues, and transferring tumor cells to distant tissues, all lead to tumor development and progressions (9). This only provides non-surgical treatment options for patients and therefore, opting for targeted therapy that entails precision and selective killing mechanisms to reduce damage to healthy tissues must be adapted using potential biomarkers for tumor treatment.

Cyclin-dependent kinase regulatory subunit 1B (CKS1B), a member of the conserved cyclin kinase subunit 1 (CKS1) protein family contributes greatly to cell cycle modulation. The cell cycle regulation depends on interactions of the cyclin-dependent kinase (CDK) and its inhibitors. The CKS1B protein is encoded by the *CKS1B* gene present on the human chromosome Iq21, has a molecular weight of 9 kDa, and consists of about 79 amino acids. *CKS1B* regulates the eukaryotic mitotic cycle and play important role in normal cell division and growth. Moreover, functional analysis represented a significant impact of CKSIB expression in the cell division cycle (10). Dysregulations of the cell cycle, as well as impaired cell differentiation as a consequence of CKS1B expression changes, can trigger malignant tumor progressions. CKS1B binds and regulates the functions of cyclin-dependent protein kinase catalytic subunits (11, 12), and is shown to be involved in the promotion of cell growth, invasion, metastasis, and chemical resistance (13, 14). Overexpression of the *CKS1B* gene has also been evident in multiple cancers including - hepatocellular carcinoma (13), colon cancer (15), lung cancer (16), oral squamous cell carcinoma (17), breast cancer (18), and retinoblastoma (RB) (12), among others. The *CKS1B* gene has also been associated and identified as one of the 70 high-risk genes, the expression of which is inversely proportional to the survival of patients newly diagnosed with multiple myeloma (MM) (19); its high nuclear expression is a poor prognostic factor in relapsed/refractory MM patients (20). These findings coupled with additional database analyses indicate the involvement of *CKS1B* as a therapeutic candidate (21, 22).

Despite the high prevalence of LUAD and LUSC among NSCLCs, treatment options and survival rates become limited due to a lack of early detection. Prognostic factors help identify patient characteristics as well as the stage of cancer, before the treatment. Some of these factors for patient survival in the LC subtypes include - performance status, stage-tumor dimension, nodal status, and weight loss. The absence of initial symptoms, as well as the inefficiencies of CT scans or chest radiographs to detect cancers in their early stages, are the contributors to the delayed treatment response and the low survival rates. Targeted therapy utilizing biomarkers is a field that holds great prognostic value but is yet unavailable for LC detection due to inadequacies in robust sensitivity and specificity of the biomarkers or their functional relevance with lung carcinogenesis (23). Therefore, in our study, we upheld the hypothesized significance and importance of the *CKS1B* expression as a potential biomarker for early diagnosis and treatment of LUAD and LUSC, exhibiting its immunotherapeutic features and prognostic value.

## 2. Materials and Methods

### 2.1. Determination of *CKS1B* mRNA expression in cancerous and normal tissues

Three different databases are entitled Oncomine (https://www.oncomine.org) (Rhodes et al., 2004), GEPIA2 (http://gepia2.cancer-pku.cn) (24), and GENT2 (http://gent2.appex.kr) (25) were used to analyze the expression of *CKS1B* expression in multiple cancerous tissues and their normal, non-cancerous counterparts. The Oncomine database contains 715 independent datasets that store experimental information of 86,733 cancer samples and 12,764 standardized tissue samples. This server is well-known for its high reliability, accuracy, consistency, and scalability (26). In the Oncomine database, the default parameters i.e., p-value threshold of 1E-4, fold change threshold of 2.0, and Top 10% Gene rank, were set. GEPIA2 (Gene Expression Profiling Interactive Analysis) database is widely utilized to analyze the mRNA expression profiling of a variety of genes. The server houses the information of 9,736 cancerous and 8,587 normal tissue samples, extracted from the TCGA (The Cancer Genome Atlas) and GTEx (Genotype-Tissue Expression) projects (24). During analysis by the GEPIA2 server, all the parameters were kept at their default values. Along with the above-mentioned servers, GENT2 (Gene Expression database of Normal and Tumor tissues-2) was also used to analyze the expression of the *CKS1B* gene in more than 68,000 tumor and normal tissue samples stored in its database. This server generates reliable and quite accurate outcomes by exploiting the Apache Lucene indexing and Google Web Toolkit (GWT) framework (25). Finally, the expression of the *CKS1B* gene was observed for both LUAD and LUSC, according to the TIMER analysis (http://timer.cistrome.org/).

### 2.1. Expression profiling of *CKS1B* in cancerous and normal lung tissues

Four different web servers namely Oncomine (26), GEPIA2 (24), UALCAN (http://ualcan.path.uab.edu/) (27), and HPA (https://www.proteinatlas.org/) (28) were used to determine the expression profile of *CKS1B* mRNA in normal and cancerous LUAD and LUSC tissues. UALCAN web-server allows the users to detect novel cancer biomarkers and carry out numerous *in silico* analyses to map the expression of target genes by providing free access to the cancer OMICS datasets like the TCGA database (27). The relative expression pattern of *CKS1B* mRNA in both LUAD and LUSC samples was also analyzed using the TCGA database. The HPA (Human Protein Atlas) database was used to make a visual comparison between LUAD and LUSC tissues and normal lung tissue. This project aims to map all human proteins in cells, tissues, and organs by combining various omics technologies such as mass spectrometry-based proteomics, transcriptomics, antibody-based imaging, and so on (28). The default parameters for all these servers were used and the p-value less than 0.05 was considered significant.

### 2.2. Correlation of *CKS1B* overexpression with clinical features and promoter methylation

The UALCAN server was used to determine the correlation of *CKS1B* overexpression with clinical features (27). Individual cancer stages, patient’s race, patient’s gender, patient’s age, patient’s smoking habit, tumor histology, nodal metastasis status, and T53 mutation status are the clinical features that were analyzed in the server keeping all the parameters default. After that, the UCSC Xena Functional Genomic Explorer (https://xenabrowser.net/) was used to find out the DNA methylation pattern of *CKS1B* gene promoter associated with LUAD and LUSC (29). The TCGA server analyzed experimental data of 706 LUAD and 626 LUSC samples, and the DNA Methylation of 27k and 450k patterns of *CKS1B* for LUAD and LUSC respectively.

### 2.3. Analyses for mutations and copy number alterations

The pattern of mutations and copy number alterations of the *CKS1B* gene in association with LUAD and LUSC was determined using the cBioPortal server (https://www.cbioportal.org/; designed to explore, visualize, and interpret multidimensional cancer genomics data) (30). In this study, a total of 2,983 and 1,176 LUAD and LUSC samples respectively, were exploited. The mutation spectrum of the *CKS1B* gene responsible for cancer development was determined by the cBioPortal server, keeping all the parameters as default.

### 2.4. Analysis of correlation between *CKS1B* expression and the survival of LC patients

PrognoScan server (http://dna00.bio.kyutech.ac.jp/PrognoScan/) was exploited to determine the relationship of the survival of patients with LUAD and LUSC and *CKS1B* gene expression. PrognoScan is a database for meta-analysis of genes that scans the freely available microarray datasets to predict the interaction between the expression of a query gene and the prognosis of cancer (31). The server uses the Kaplan-Meier statistical method to plot the gene expression in the X-axis and the possibility of patient survival in the Y-axis. During the analysis, default parameters were used and any cox p-value less than 0.05 was considered to be statistically significant.

### 2.5. Analysis of the genes co-expressed with *CKS1B* in LC tissues

Three different servers i.e., Oncomine (26), GEPIA2 (24), and UCSC Xena web browser (29) were used to determine the genes which tend to co-express with *CKS1B*. Firstly, the Oncomine server was used to search the list of co-expressed genes. The server ranks the co-expressed genes based on correlation scores. Thereafter, the analysis of the relationship between the *CKS1B* gene and the gene that generated the highest correlation score in the Oncomine server was carried out using the GEPIA2 server. Finally, the UCSC Xena web browser was utilized to design the gene expression pattern of the selected genes in LUAD and LUSC patients.

### 2.6. Network and Pathway Analysis in LUAD and LUSC for the *CKS1B* Gene

GeneMANIA (https://genemania.org) is a web-based platform that uses a large database of functional interaction data to assess the relationship between a gene of interest and other genes. Centered on protein and genetic interactions, pathways, co-expression, co-localization, and protein domain similarity, this platform was used to examine the relationship of the *CKS1B* gene with other genes. In addition, using the International Molecular Exchange Consortium (IMEx) protein interactions database, the previously reported 20 associated genes and *CKS1B* was used in NetworkAnalyst (https://www.networkanalyst.ca/) to establish the protein-protein interaction at a generic level. The RTK/Ras pathway, P13K/AKT pathway, MAPK, and Rap1 signaling pathway were found to involve *CKS1B* in the KEGG 2019 database. Following that, PathwayMapper in the cBioPortal server was used to analyze these pathways in detail as well as to estimate the frequency of *CKS1B* alterations.

### 2.7. Determination of *CKS1B* ontology and signaling pathways of cancer-associated genes

The Enrichr server (http://amp.pharm.mssm.edu/Enrichr/) was used for gene ontology (GO) and cell signaling pathways of *CKS1B*. The server generates the annotation enrichment results of a target gene set by comparing multiple genomics datasets containing previous biological knowledge (32, 33). To do the analysis, the *CKS1B* along with the other co-expressed genes found in the Oncomine server was used. The Go terms (i.e., GO biological process, GO molecular function, and GO cellular component) and the signaling pathways (from BioPlanet 2019, KEGG 2019 Human, and Reactome 2016 databases) were analyzed using the Enrichr web server.

## 3. Results

### 3.1. Expression of *CKS1B* mRNA in cancerous and non-cancerous tissues

*CKS1B* is substantially upregulated in various cancer cells, including LC, according to 56 studies out of 452 retrieved datasets (**Figure 1A**). GEPIA2 results also indicate a similar pattern of *CKS1B* upregulation in 33 different types of cancer cells including LUAD and LUSC (**Figure 1B**). *CKS1B* is overexpressed in various cancers, including LUAD and LUSC, according to a review of GENT2 databases using HG-U133pLUS 2 and HG-U133A platforms (**Figure 1C**). In addition, the TIMER analysis revealed that *CKS1B* was overexpressed in both LUAD and LUSC as compared to other cancer forms (**Figure 1D**). Overall, the findings showed that the *CKS1B* gene is significantly upregulated in LC tissues as compared to normal tissues and other cancer cells (**Figure 1**).

**Figure 1.**
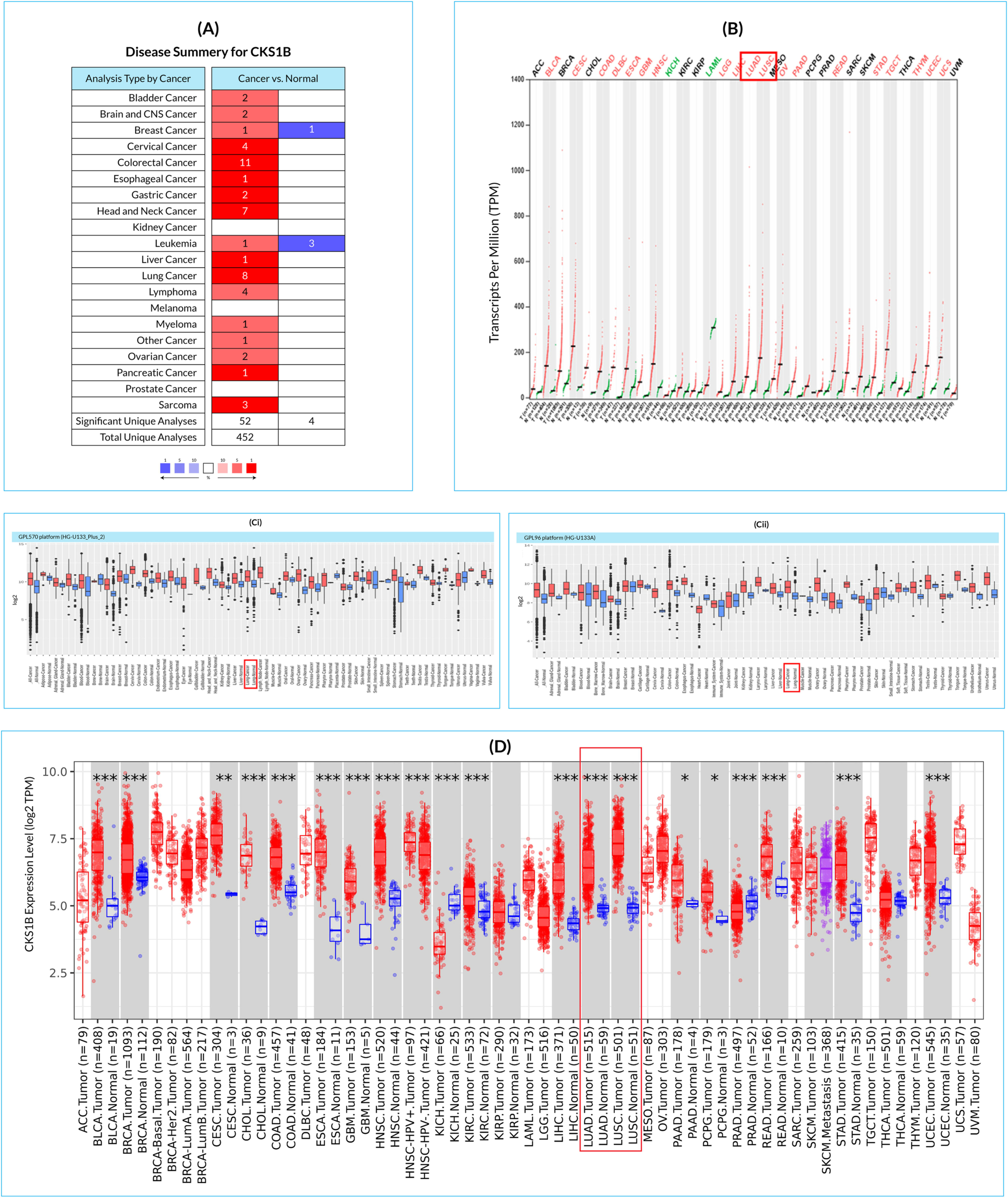
*CKS1B* tissue-wide expression profile in multiple cancer types. (A) Contrast of cancer versus healthy tissues, with high and low mRNA expression shown by red and blue colors, respectively, (B) the dot plot illustrates the gene expression profile of the *CKS1B* gene in 33 diverse human cancers, including tumors and healthy tissue samples combined. (C) the box-plot representing the *CKS1B* mRNA expression in cancers and corresponding healthy tissues by using HG-U133Plus 2 (Ci) and HG-U133A platforms (Cii) of the GENT2 database, where the boxes represent the median, the dots denote the outliers, and the red-boxes correspond to cancerous tissue, and the blue-boxes refer to the expression of healthy tissues. (D) *CKS1B* mRNA expression in LUAD and LUSC tissue relative to normal tissue (statistical significance measured using differential analysis, *P 0.05; **P 0.01; ***P 0.001).

### 3.1. Expression of *CKS1B* transcript in human LC tissues

Comparative analysis using Oncomine server between LC (LUAD and LUSC) and normal tissue revealed an increased expression of the *CKSIB* gene in both LC subtypes (**Figure 2Ai-vi & Supplementary Table S1**). Additional evaluation of TCGA datasets using the UALCAN and GEPIA2 server also represented a significant overexpression of *CKS1B* in the LC subtypes (**Figure 2Bi, 2Bii, and 2C**). Moreover, relative immunohistochemistry analysis of healthy and cancerous tissues using the HPA database showed moderate to weak staining signals in normal alveolar cells (**Figure 2F**), and strong signals in both LUAD (**Figure 2D**) and LUSC (**Figure 2E**) tissue samples (**Figure 2 and Supplementary Table S1)**.

**Figure 2.**
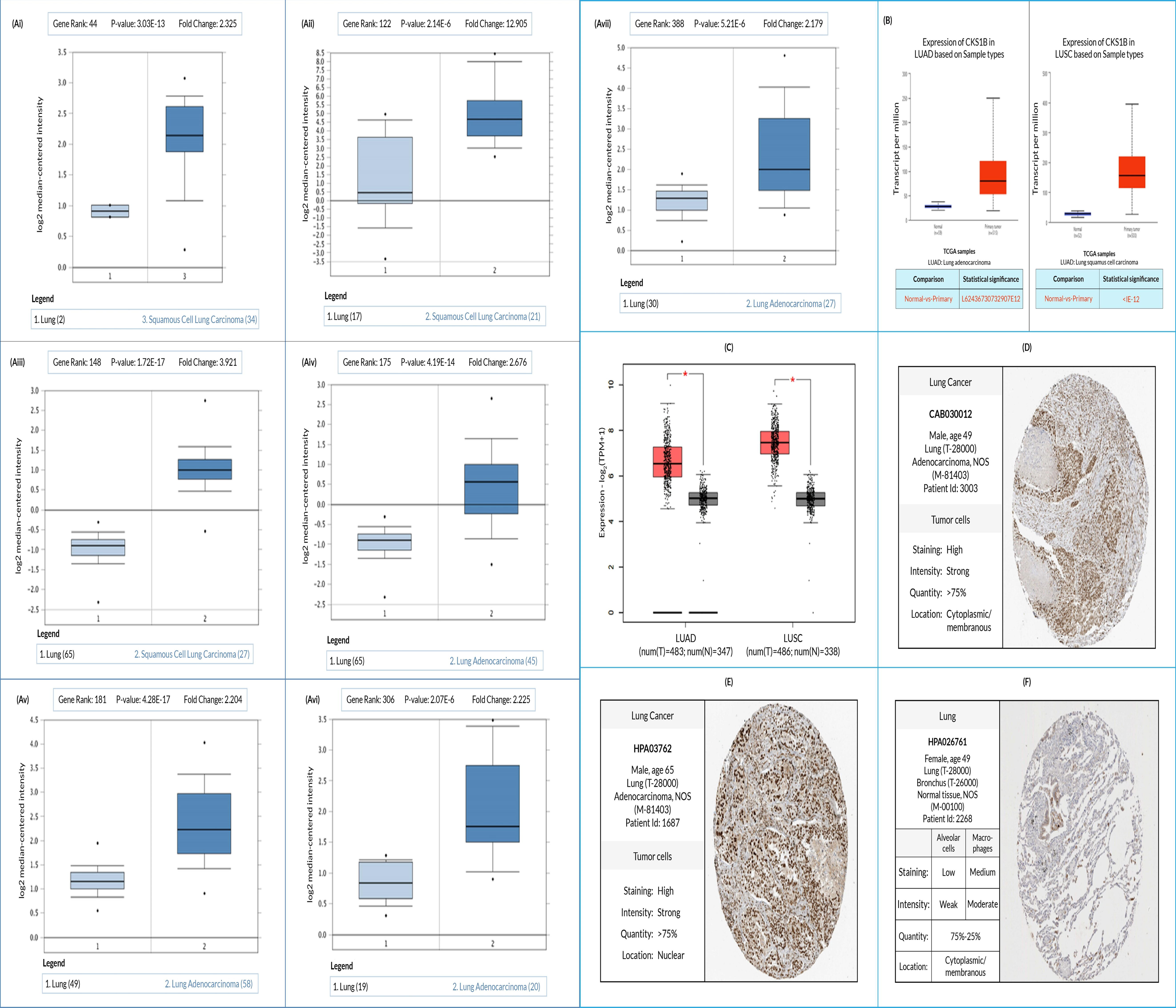
Evaluation of *CKS1B* expression in LUAD and LUSC, with (A) box-plots showing comparative expression between normal (left) and cancer tissue (right) - for LUAD and LUSC (Ai-Avii), (B–C) box-plots showing *CKS1B* mRNA expression dependent on sample forms in tumor tissue and normal tissues for LUAD (Bi) and LUSC (Bii), using the UALCAN and GEPIA2 (C), respectively, and (D-F) the immunohistochemistry images of *CKS1B* expression in LUAD (D) and LUSC (E) tissues as well as normal tissue (F) extracted from the HPA database.

### 3.2. Correlation between *CKS1B* expression and clinical features of LC patients

Analysis of the UALCAN database revealed that *CKS1B* upregulation for LUAD and LUSC varied in individual cancer stages (**Figure 3Ai, ii**) due to patient ethnicity, sex, age, smoke addiction, tumor histology, nodal metastasis, and TP53 mutation status (**Figure 3, Supplementary Table S2 & S3**). Correlated overexpression of *CKS1B* was also found in the patient of Asian (**Figure 3Bi, ii**), male (**Figure 3Ci, ii**), age group range of 21-40 years old (**Figure 3Di and ii**), smoking habit (**Figure 3Ei, ii**), tumor histology (**Figure 3Fi, ii**), nodal metastasis (**Figure 3Gi, ii**), and TP53 mutation status (**Figure 3Hi, ii**), for both LUAD and LUSC **(Supplementary Table S2 & S3**). These findings suggest a potential correlation of *CKS1B* upregulation and clinical characteristics with LC than normal patients.

**Figure 3.**
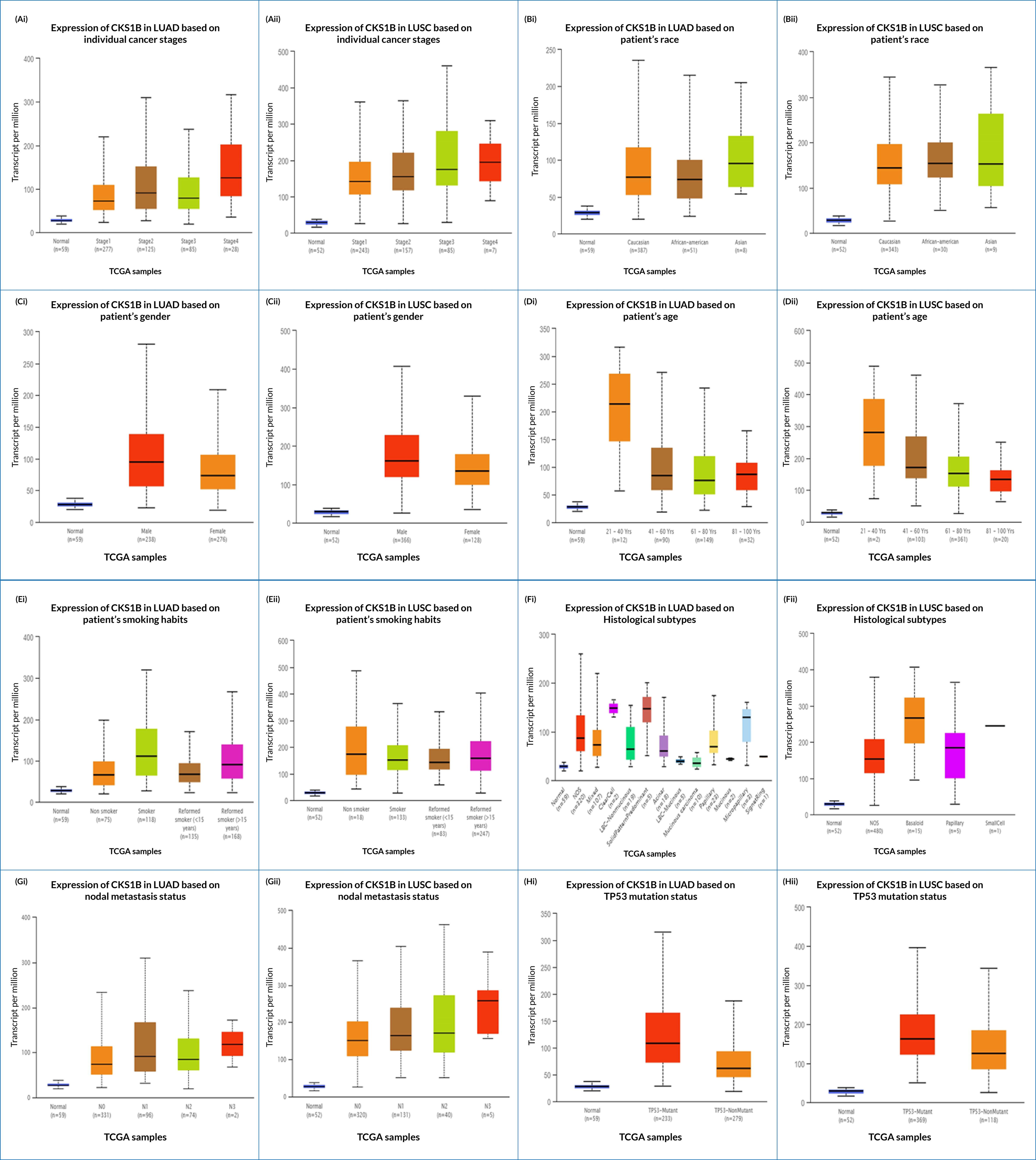
*CKS1B* expression and diagnostic criteria correlation in LUAD and LUSC patients. Individual cancer stage, patient’s ethnicity, sex, age group, smoke addiction, histological subclasses, nodal metastasis, and TP53 status are seen in the *CKS1B* mRNA expression in LUAD (Ai-Hi) and LUSC (Aii-Hii), respectively.

### 3.3. TCGA dataset utilization for analysis of LC promoter methylation

Association between *CKS1B* expression and DNA methylation was evaluated using two separate methylation patterns present in the server, namely Human Methylation 27k and Human Methylation 450k for both the LC subtypes (**Figure 4A, 4B)**. Retrieved heat maps showed a close negative correlation between *CKS1B* expression and some CpG islands in LUAD (**Figure 4A**), whereas LUSC (**Figure 4B**) suggested that the overexpression of *CKS1B* associated with decreased DNA methylation in LUAD and LUSC, and vice versa (**Figure 4**).

**Figure 4.**
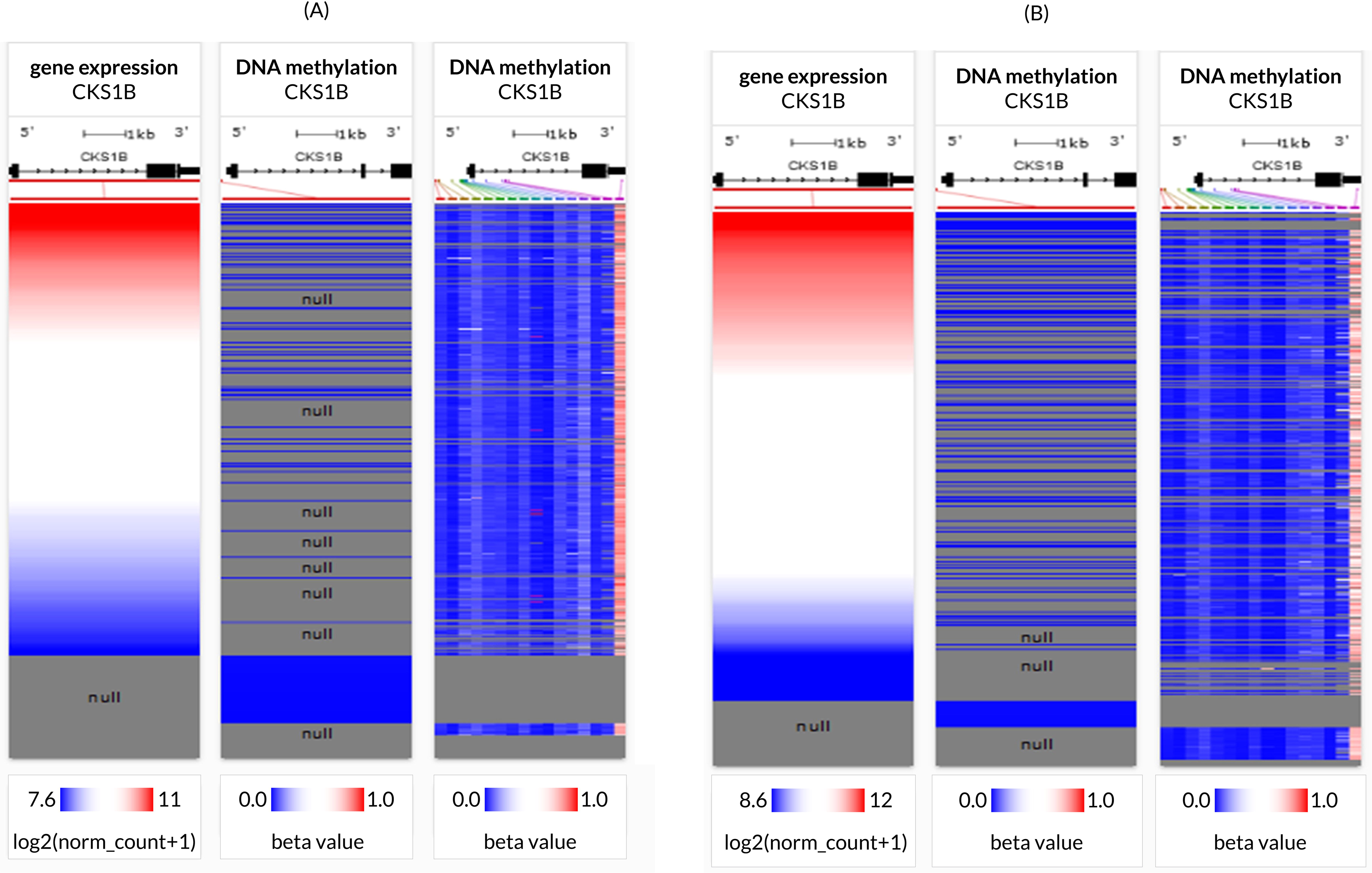
*CKS1B* gene promoter methylation in LC tissues. For LUAD (A) and LUSC (B), a heat map of *CKS1B* expression and DNA methylation status was generated.

### 3.4. Analyzing mutations, copy number alterations, and expression of mutant *CKS1B* transcripts

An analysis of the *CKS1B* gene alterations in both the subtypes of LC was done using the cBioPortal database; totaling 2983 samples from 8 LUAD studies and 1176 samples from 3 LUSC studies (**Table 1**). Studies revealed 191 (6%) alteration in the *CKS1B* gene in LUAD samples (**Figure 5Ai**), with a somatic mutation frequency of 0.1 percent and 42 (4%) altered *CKS1B* gene in LUSC samples with a somatic mutation frequency of 0.3 percent (**Figure 5Aii**). *CKS1B* is found between 154974681 and 154979251 bp on chromosome 1q21.3. (http://atlasgeneticsoncology.org/Genes/CKS1BID40092ch1q21.html). Two mutations including one missense and one nonsense were found between 40-50 residues of the CKS domain of the 79 AA long CKS1B protein in LUAD samples (**Figure 5Ai**) whereas three missense mutations in the same mutation site (7th residue), as well as one fusion, were found in LUSC samples (**Figure 5Aii**). Furthermore, as seen in **Figure 5Bi**, amplification was found to be the most prevalent alteration type in both LC subtypes, with the highest frequency of 14.78 % of 230 cases in the LUAD TCGA, Nature 2014 dataset. The LUSC TCGA, Firehose Legacy dataset had the highest alteration frequency of the three, at 4.18 % of 502 cases. In the alteration study of LUAD TCGA, Nature 2014 dataset, both mutation and amplification of *CKS1B* have been identified with a frequency of 0.2% (1 case) and 3.98% (20 cases), respectively. In the LUSC TCGA, PanCancer Atlas, a fusion was also observed with a frequency of 0.21 % (**Figure 5Bii**). Additionally, the *CKS1B* mRNA expression profile exhibited higher amplification, fewer gains, and twelve cases of shallow deletions in LUAD samples (**Figure 5Ci**). In LUSC samples, two missense mutations and one fusion along with some gains, diploids, and shallow deletions were found, as appeared in **Figure 5Cii**.

**Table 1.**
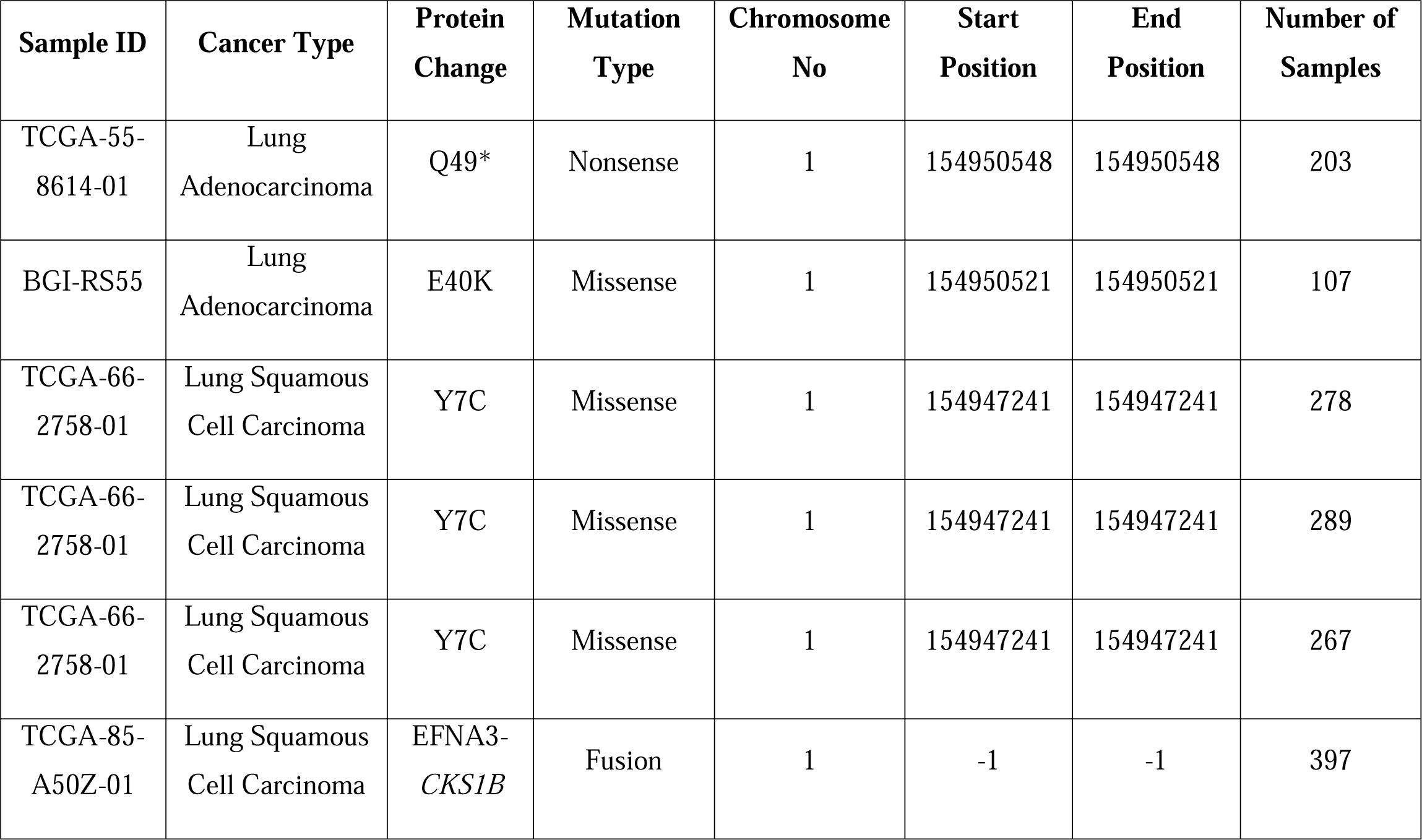
Selected *CKS1B* mutational positions and types in LUAD and LUSC from the TCGA dataset.

| Sample ID | Cancer Type | Protein Change | Mutation Type | Chromosome No | Start Position | End Position | Number of Samples |
| --- | --- | --- | --- | --- | --- | --- | --- |
| TCGA-55-8614-01 | Lung Adenocarcinoma | Q49* | Nonsense | 1 | 154950548 | 154950548 | 203 |
| BGI-RS55 | Lung Adenocarcinoma | E40K | Missense | 1 | 154950521 | 154950521 | 107 |
| TCGA-66-2758-01 | Lung Squamous Cell Carcinoma | Y7C | Missense | 1 | 154947241 | 154947241 | 278 |
| TCGA-66-2758-01 | Lung Squamous Cell Carcinoma | Y7C | Missense | 1 | 154947241 | 154947241 | 289 |
| TCGA-66-2758-01 | Lung Squamous Cell Carcinoma | Y7C | Missense | 1 | 154947241 | 154947241 | 267 |
| TCGA-85-A50Z-01 | Lung Squamous Cell Carcinoma | EFNA3-*CKS1B* | Fusion | 1 | -1 | -1 | 397 |

Consequently, the observations suggest that *CKS1B* overexpression in LUSC and LUAD may not have an established correlation with mutations or copy number alterations in the *CKS1B* gene (**Figure 5**). Nevertheless, these genetic modifications can serve as prognostic measures for LUAD and LUSC, using *CKS1B* as a biomarker. Also, the correlation of *CKS1B* to other functional proteins which are important for cell cycle, proliferation, and DNA repair, makes it a key component for increasing human susceptibility to diseases or detrimental effects, in case of mutations or alterations. This raises the importance of thorough future evaluations in regards to mutations and copy number alterations.

**Figure 5.**
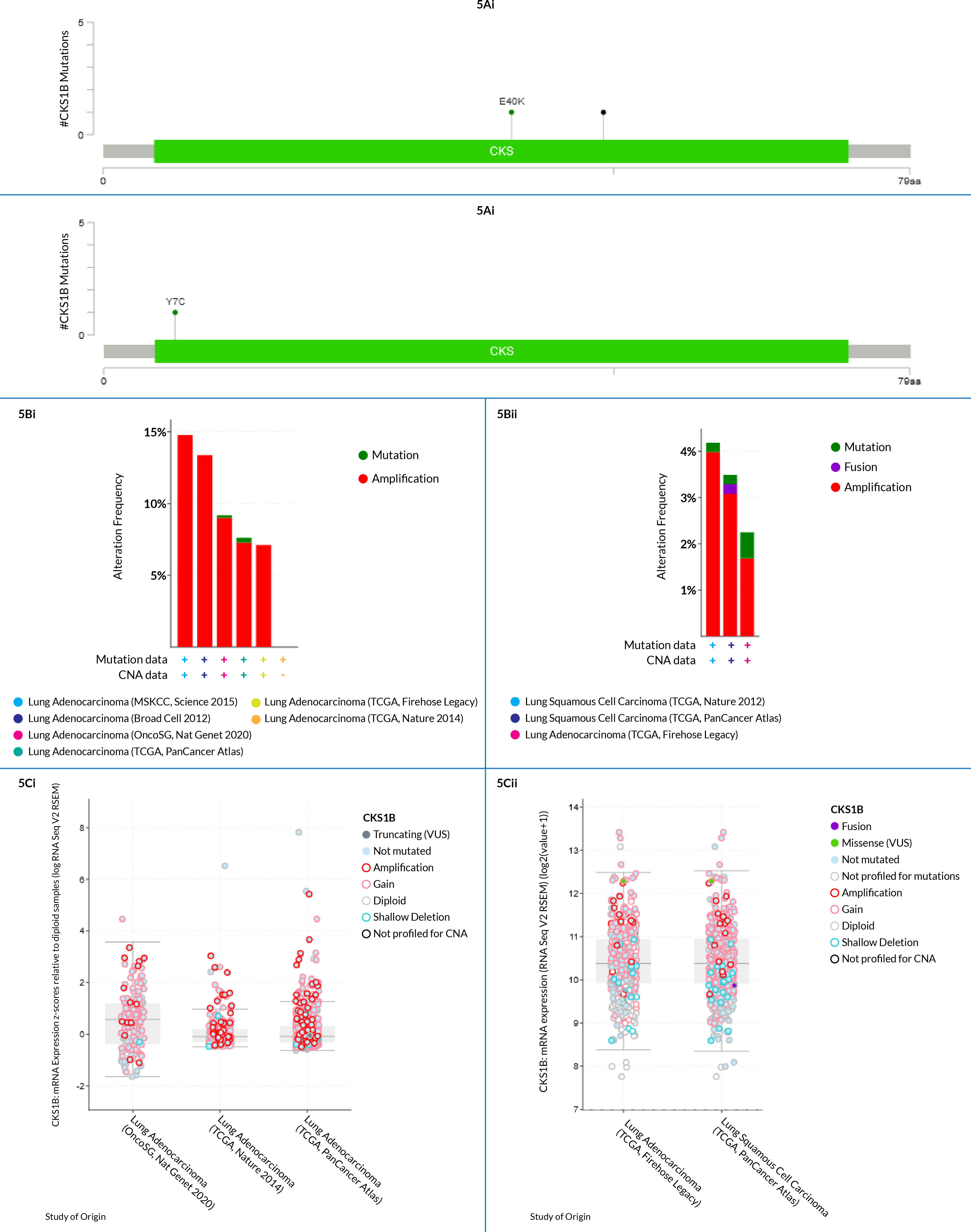
*CKS1B* gene alterations and mutations in LUAD and LUSC tissues. (Ai) lollipop plot shows the type of alteration in two mutation spots within the peptide sequence (40-50 residues) of CKS domain in *CKS1B* from LUAD tissues, (Aii) represents the alterations in a single mutation spot in *CKS1B* from LUSC tissues, (Bi and Bii) the bar diagrams indicate the mutation frequencies and genome alterations in the *CKS1B* gene for LUAD and LUSC, respectively, and (Ci and Cii) indicate the correlation between the expression and copy number alterations of *CKS1B* for LUAD and LUSC in the TCGA dataset

### 3.5. Correlation between *CKS1B* expression and clinical prognosis of LC patients

A negative correlation between *CKS1B* expression and LC patient survival (significant level was held at Cox p-value 0.05 and HR>1) was found using the PrognoScan database. *CKS1B* expression was observed to be negatively associated with patient survival. This finding indicated that patients with elevated *CKS1B* expression might have a lower survival risk, whereas those with poor or average *CKS1B* expression may have a higher survival (overall or relapse-free) rate (**Figure 6** & **Supplementary Table S4)**. The datasets showed an increased overall survival risk in patients with low *CKS1B* expression, while those with higher *CKS1B* expression had a considerably low overall survival rate (**Figures 6**). The investigation suggests a correlation between increased *CKS1B* expression and a poor prognosis in LC patients.

**Figure 6.**
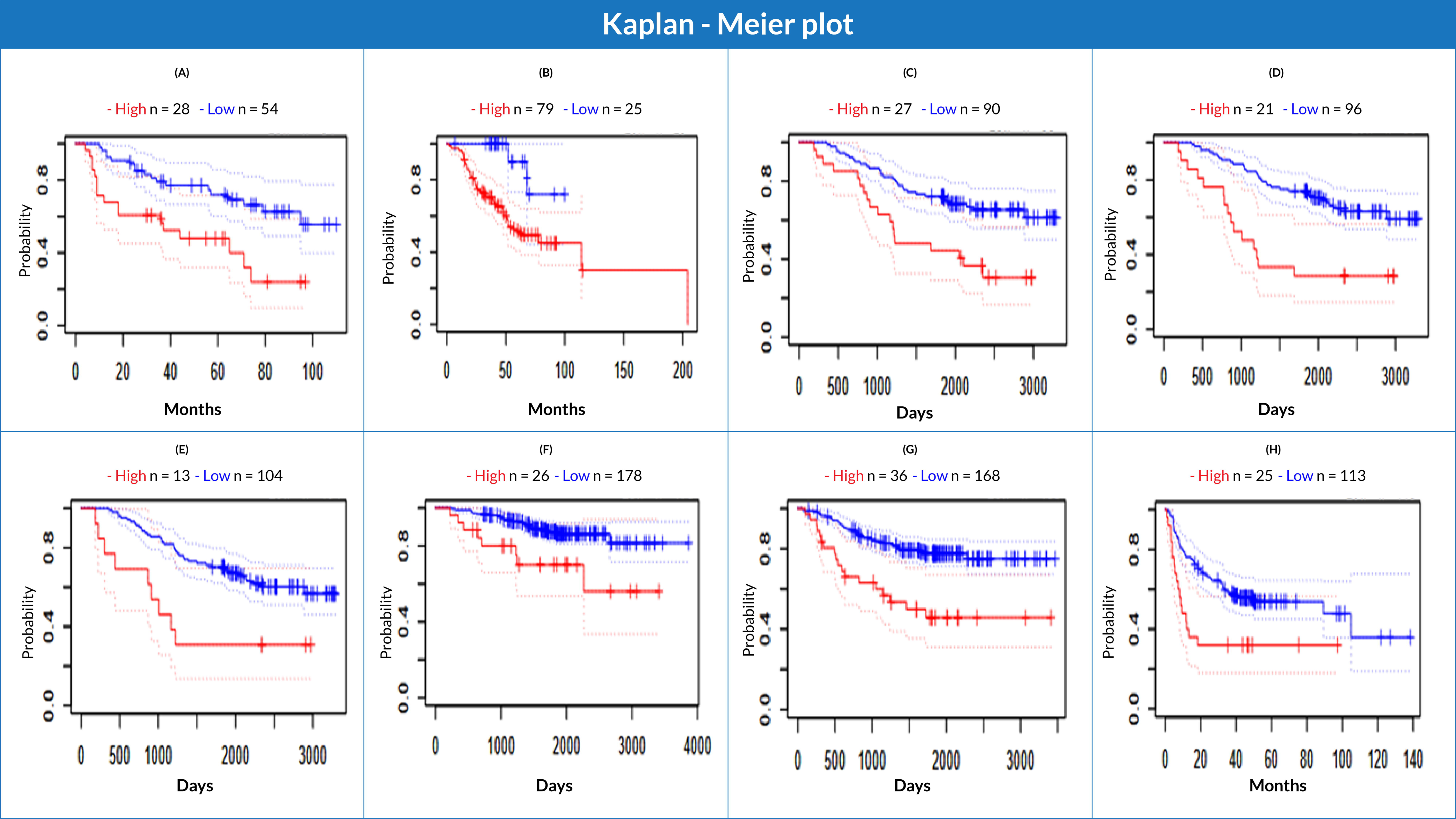
The correlation between *CKS1B* gene expression and LC patient survival is represented by the Kaplan-Meier plot. The survival curves show patients’ survival in plots of high (red) and low (blue) expression of *CKS1B*, with A–F indicating overall survival and G, H indicating relapse-free survival. The study emphasized on the expression of *CKS1B* in LC patients.

### 3.6. Determination of *CKS1B* and human LC associated gene signatures

The Oncomine database was utilized to investigate the co-expression profile of *CKS1B* with 20 other genes, using 34 samples from LUAD and LUSC patients (**Figure 7A**). Among the 20 genes analyzed, the most co-expressed (R = 1.00) was the Src homology 2 domain-containing transforming protein C1 (SHC1). Following this, the GEPIA2 server also indicated a positive correlation with a Spearman coefficient in both LUAD (R = 0.25) and LUSC (R = 0.21) between *CKS1B* and *SHC1* (**Figure 7Bii**). This was further supported by Pearson and Spearman correlation analyses results obtained from the UCSC Xena server, using the TCGA database (**Figure 7C-D**). The Pearson correlation values for LUAD (**Figure 7Ci-Di**) and LUSC (**Figure 7Cii-Dii**) patients were 0.3563 and 0.2327, and the Spearman correlation values for LUAD and LUSC were 0.4073 and 0.2176, respectively (**Figure 7C-D**). These values suggest that *CKS1B* and *SHC1* can share a common biosynthetic pathway, or combine to form a large protein complex with LC progression.

**Figure 7.**
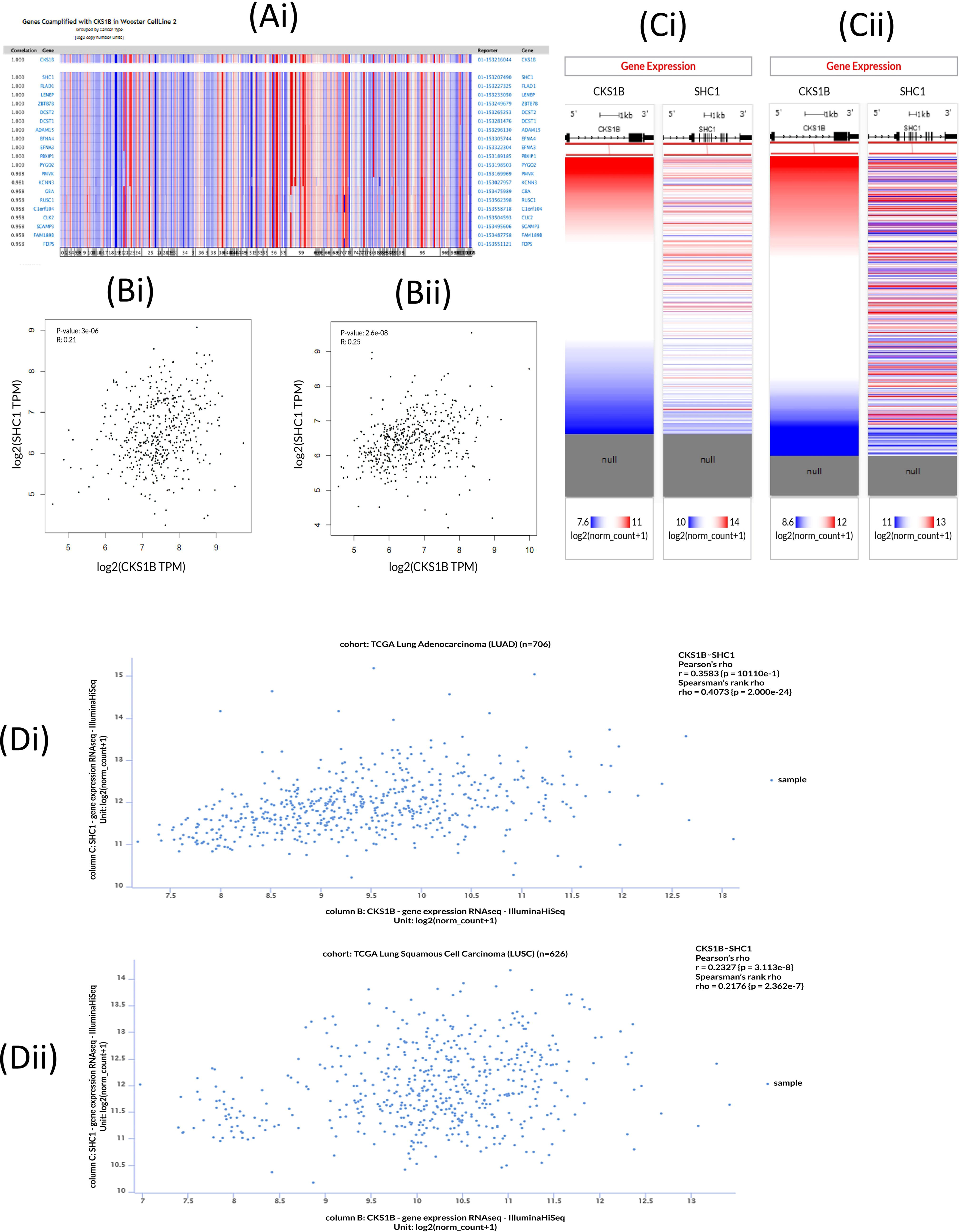
*CKS1B* and co-expressed gene profile in human LC. (A) The co-expression profile of *CKS1B* retrieved from the Oncomine database, (B) *CKS1B* and *SHC1* correlation analysis obtained from the GEPIA2 server, (C) *CKS1B* and SHC1 mRNA expression heatmap from the TCGA database, and (D) the co-expression analysis between *CKS1B* and *SHC1* genes in LC using the UCSC Xena server.

### 3.7. Analysis of *CKS1B* gene network in LUAD and LUSC

The GeneMANIA server retrieves a complete network of *CKS1B* gene and its crosstalk genes of interaction in LC, displaying the physical interactions (67.40%), co-expression (13.82%), predicted (6.33%), co-localization (6.14%), pathways (4.33%), genetic interactions (1.39%) and shared protein domains (0.59%) (**Figure 8A**). Moreover, based on the International Molecular Exchange Consortium (IMEx) protein interactions database, protein-protein interaction (PPI) network was constructed for the co-expressed genes, and using NetworkAnalyst, these interactions were further analyzed for LUAD and LUSC progressions. **Figure 8B** shows the PPI network in which, the degree of a node is the number of connections among the node, and betweenness is the smallest path amongst nodes showing *SHC1* (Degree:207, Betweenness:65895.06), *MUC1* (Degree:31, Betweenness:7217.707), *ADAM15* (Degree:31, Betweenness:9000.452), *TRIM46* (Degree:30, Betweenness:10467.46), *DAP3* (Degree:28, Betweenness:8994.579), *CKS1B* (Degree: 27, Betweenness: 9277.062), and *ZBTB7B* (Degree: 19, Betweenness: 6702.692) as the major proteins of the network.

**Figure 8.**
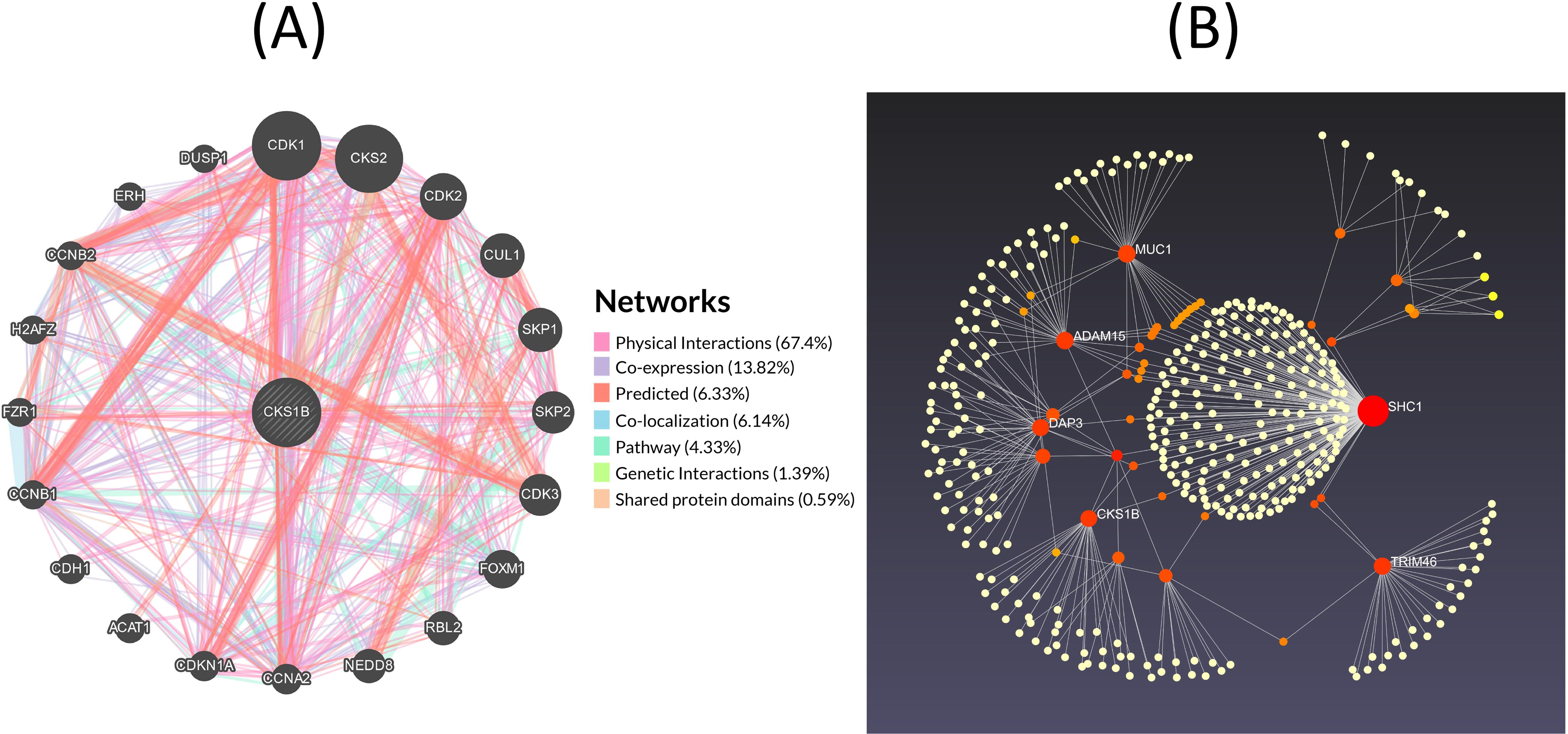
Analysis of gene networks. (A) *CKS1B* gene with its neighboring genes physical interactions (67.40%), co-expression (13.82%), predicted (6.33%), co-localization (6.14%), pathways (4.33%), genetic interactions (1.39%) and shared protein domains (0.59%) (B) Protein-protein interaction network represented by IMEx protein interactions database.

### 3.8. Evaluation of *CKS1B* gene associated pathways in LUAD and LUSC

The PathwayMapper tab in the cBioPortal server exhibits the *CKS1B* alteration frequency (in percentage) over multiple pathways on the LUAD and LUSC datasets (**Figures 9A–B**). RTK-Ras-RAF, PI3K/AKT, and TP3 pathways are mainly altered due to *CKS1B* gene products. These alterations coordinate the progression of LUAD and LUSC. Alterations related to *CKS1B* mainly induce the changes of EGFR (18.8%), KRAS (20.5%), FGFR1 (8.5%), and BRAF (6.0%) genes for regulation of RTK-Ras signaling pathway (**Figure 9A**) as well as PTEN (6.4%), PIK3CA (18.0%), STK11 (10.8%), and RICTOR (9.8%) in the regulation of P13K/AKT signaling pathway (**Figure 9B**); TP53 (48.9%), and CDKN2A (22.4%) in TP53 signaling pathway (**Figure 9C**) for cancer proliferation and development.

**Figure 9.**
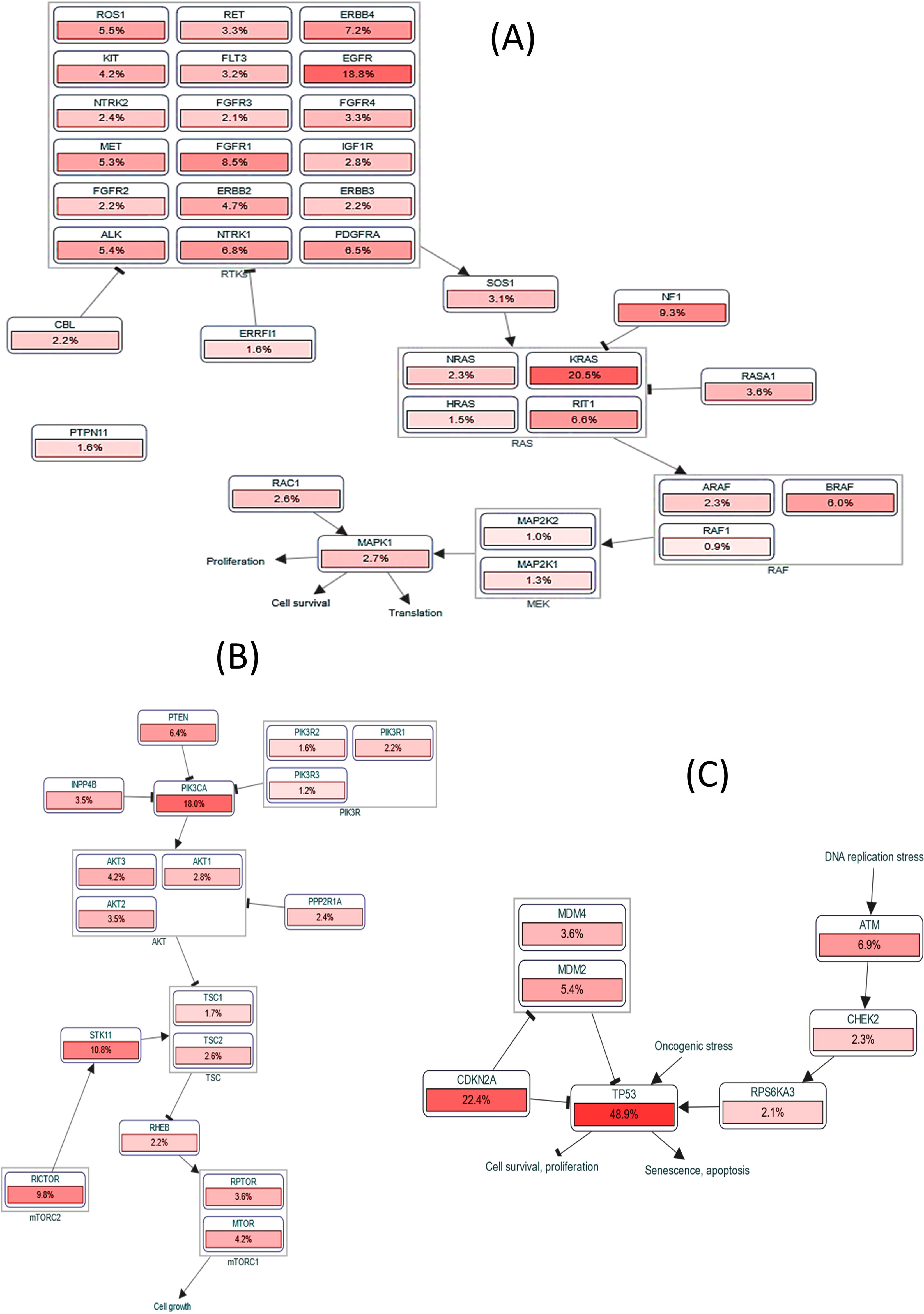
Pathway exploration. (A–C) *CKS1B* and related gene impact in modulating alteration frequency of (A) RTK-Ras-RAF signaling pathway, (B) P13K/AKT signaling pathway, (C) TP53 pathway.

### 3.9. Identification of *CKS1B* ontologies and related signaling pathways involved in LC

Results from three databases shown in **Figure 10A-C** were used for pathway determination. The KEGG human 2019 database analysis revealed different significant associations of different pathways in LC progression including RTK/Ras pathway, P13K/AKT pathway, MAPK, and Rap1 signaling pathway (**Figure 10A**). Additionally, the Reactome 2016 database showed LC progression depends on the pathways related to EPHA or EPH-Ephrin mediated signaling and repulsion of the cells, etc. (**Figure 10B**). The analysis of the Bioplanet 2019 database also revealed the Ephrin receptor A forward pathway, and p27 phosphorylation regulation during cell cycle progression alongside other important pathways associated with LC (**Figure 10C**). Following this, GO terms were also checked for the corresponding genes which mainly include, positive regulation of aspartic-type peptidase activity, axon guidance and axogenesis (**Figure 10D**), ephrin receptor binding and transmembrane-ephrine receptor activity (**Figure 8E**), and possible interactions with beta-catenin-TCF complex, mitochondrial small ribosomal subunit, and an anchored component of the plasma membrane (**Figure 10F**).

**Figure 10.**
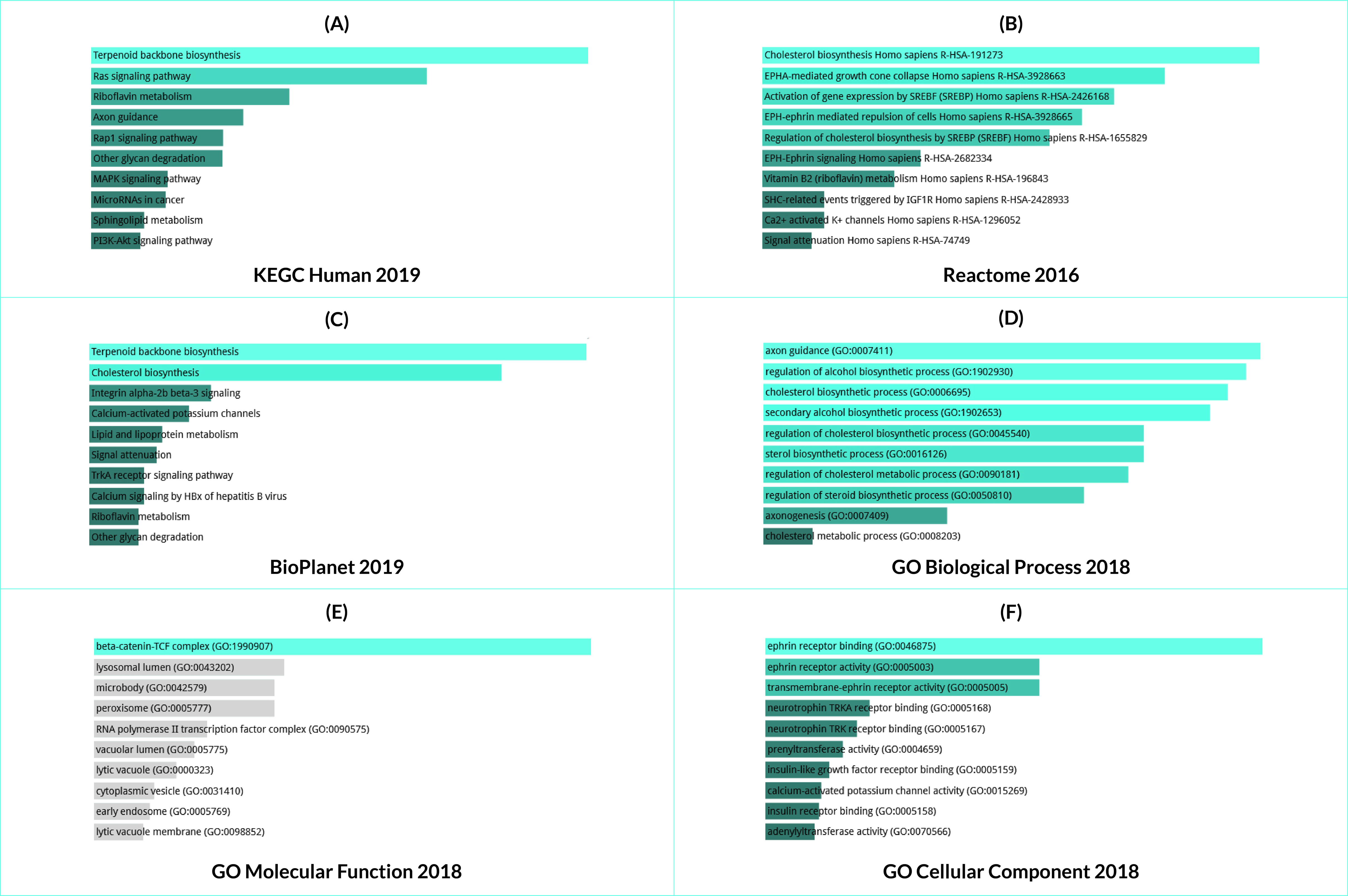
Evaluating the gene ontologies and signaling pathways associated with *CKS1B* expression and LC. Ontologies and pathways are retrieved from (A) KEGG human 2019, (B) Reactome 2016, (C) BioPlanet 2019, (D) GO biological process 2018, (E) GO molecular function GO Molecular Function 2018 GO Cellular Component 2018, and (F) GO cellular component 2018. The length is representing the significance level whereas a brighter color of the bar indicates the more significant term, and vice versa.

## 4. Discussion

Over the years, LC has persistently maintained its top rank in terms of the worldwide mortality rate (34). In contrast, NSCLC consisting of both LUAD and LUSC solely constitutes an estimated 85% of all LC cases. LC is infamously known for its frequent diagnosis of the tumor in the malignant form (35, 36). Therefore, correct prognosis and early detection are deemed crucial for a thorough understanding of the patient’s condition as well as to coordinate further treatment approaches (37, 38). Collective reports also revealed that most of the LC (approx. 75%) patients were diagnosed only after advanced stages (stage III/IV), emphasizing the need for early diagnosis (39). Additionally, NSCLC surgical resection provides a favorable prognosis in the case of small, localized tumors (stage I) with 5-year survival rates of 70–90% (40–42). This is indicative of a strong correlation between the early diagnosis of LC and increased survival rates; a factor to be considered for optimal outcomes despite the contemporary advancements and breakthroughs in effective treatment (43).

In this study, we have evaluated *CKS1B* as a prognostic marker for the early determination of LUAD and LUSC using a bioinformatics approach. The analysis led to an existing positive correlation between *CKS1B* expression levels and progression of LC. However, the *CKS1B* expression in LUAD and LUSC exhibited a negative correlation with all cases of overall and relapse-free survival, with the overall HR > 1. Moreover, it was also found that high *CKS1B* levels had an adverse effect on survival rates (**Figure 6**); this was supported by a previous study relating poor prognosis of LC patients due to increased expression of *CKS1B* (44). Apart from expression levels, the *CKS1B* expression patterns in the two LC subtypes seemed to be majorly analogous with the different clinical characteristics of LC patients including, tumor histology, patient’s race, gender, age, smoking habit, nodal metastasis status, etc. These features require extensive investigation, since a high expression level of *CKS1B* may indicate a risk of cancer transformation and progression (**Figure 3**). An in-depth understanding of immunohistochemical scoring can be gathered from computer-aided systems and digital imaging, combined with the methodological aspects of modern immunohistochemistry (45–47). The immunohistochemical data of *CKS1B* represented a strong nuclear immunoreactivity in every target LC cell; robust and intense staining of cancer cells showing higher *CKS1B* expression compared to normal alveolar cells in LUAD and LUSC tissues (**Figure 2).** A variation in somatically acquired genetic, epigenetic, transcriptomic, and proteomic alterations, constitutes a series of histopathological processes responsible for cancer progression; their presence in the genomic region gives rise to oncogenic effects (45). Moreover, when the copy number alterations, mutations, and mutant mRNA expressions of *CKS1B* were analyzed using the cBioPortal webserver, the *CKS1B* gene was found to be altered in 191 (6%) of quarried LUAD samples with 0.1% somatic mutation frequency, and 42 (4%) of the LUSC samples with 0.3% somatic mutation frequency. Additionally, missense mutation was found to be the highest in both LUAD and LUSC making up a total of 2983 and 1176 samples respectively (**Figure 5A-C).**

Upon investigating between DNA methylation on the CpG islands of gene promoters responsible for transcriptional silencing and *CKS1B* expression, a negative correlation was observed. This was again proved through studies regarding promoter methylations in multiple cancer types including LC (22). Further co-expression and correlation analysis revealed a positive correlation of 20 genes (R = 1.00) with the *CKS1B* gene, and *SHC1* was found to be highly associated with *CKS1B* expression (**Figure 7).** Through Ras/ERK and PI3K/AKT pathway inference, *SHC1* contributes to the development of NSCLC cell stemness and malignant progression (48). Additionally, a functional network of the interacting neighboring genes of *CKS1B* in lung cancer represented co-predicted co-localization, pathways, genetic interactions, and shared protein domains (**Figure 8).** While *CKS1B* interacts with Cyclin-dependent kinases (CDKs) and their regulatory subunits (CKSs), as well as with S-phase kinase-associated proteins (SKPs), and some highly regulated proteins, *CKS1B* overexpression can alter their normal functioning and eventually lead to malignancies. CDKs are serine/threonine kinases that modulate the cell cycle with the help of certain regulatory cyclins. CDK1 is highly essential since it promotes the G2/M and G1/S transitions, as well as G1 progression (49–51). Moreover, the upregulation of CDK1 protein is closely associated with the prognosis of varying malignant tumors (52, 53). *CKS1B* binding activates CDKs and it also interacts with SKP2 to promote the ubiquitination and proteasomal degradation of p27^Kip1^. However, *CKS1B* or *SKP2* overexpression leads to increased p27^Kip1^ turnover and cell proliferation (14). Moreover, there is an existing association between *CKS1B* and Cyclin-dependent kinase subunit 2 (CKS2), which is involved in cell cycle and cell proliferation processes and thus, *CKS1B* overexpression may dysregulate CKS2 consequently triggering LUAD and LUSC progression (54) (**Figure 8A**).

Additionally, further investigation suggests an interaction between the *CSK1B* gene and its neighboring mediators that induce downregulation of multiple biological signaling pathways. Since biologically significant co-expressed gene networks are controlled by the same transcriptional regulatory programs, functionally related or members of the same pathway or protein complex, they also exhibit identical expression patterns in specific cancers as the rise and fall of transcript levels of two co-expressed genes is simultaneous (55, 56). Betweenness centrality metric is one of the most significant indicators of network essentiality since proteins with high betweenness centrality scores may serve as a communication bridge between other proteins in the network (57). SHC1, MUC1, ADAM15, TRIM46, DAP3, CKS1B, and ZBTB7B are the most important proteins that are likely to have a connection as well as high betweenness among themselves to significantly trigger the prognosis of LUAD or LUSC (**Figure 8B).**

Further evaluation using the PathwayMapper tab of the cBioPortal website revealed the frequency of alteration of the various signaling cascades of RTK-Ras-RAF, PI3K/AKT, and TP3 pathways that consequently lead to LUAD and LUSC. The RTK/Ras/RAF/ERK pathway is majorly involved in regulating cell growth and survival; however, most LUADs include constitutive activation of mitogen-activated protein kinase (MAPK) signaling triggered through alterations in RTKs, downstream Ras/RAF/MEK cascade proteins, and their regulators (58, 59). These triggered alterations of the RTK/Ras/RAF pathway have been identified in 70%–80% of LUADs, and the genes include *EGFR*, *BRAF*, and *KRAS* (60, 61, 34). While FGFR1 focal amplification was predominantly found in LUSC (21%-22%), *KRAS* mutations were observed in both LUAD (16% to 40%) and LUSC (low frequency) (62, 63). There is also a correlation between *EGFR* and *BRAF* mutations and LUAD and LUSC (64, 65); *CKS1B* gene product may induce these alterations in the RTK/Ras/RAF pathway stimulating cancer progression (**Figure 9A).** The PI3K/AKT/mTOR pathway in LUAD and LUSC has been involved in both tumorigenesis and the progression of the disease (66). In contrast, the PI3K pathway alterations including the RTK upstream regulator of PI3K, catalytic subunit PIK3CA, PTEN negative regulator, and the downstream regulator of PI3K act as cancer triggers (67). Somatic mutations and amplification of PIK3CA along with homozygous and heterozygous deletions of PTEN correlate to tumorigenesis in LUAD and LUSC (68) (**Figure 9B**). Our study exhibits the impact of *CKS1B* gene product in the alteration of the aforementioned genes by downregulating RTK-Ras-RAF and PI3K/AKT signaling pathways (**Figure 9A-B**), as well as in the alteration of TP53 and CDKN2A through the downregulation of the TP53 pathway (**Figure 9C),** which may induce cancer progression. The ephrin-mediated signaling pathway links to *CKS1B* and the correlated genes inducing LUAD and LUSC progressions. Expression patterns of Eph receptor tyrosine kinases and their ephrin ligands in cancer cells and tumor blood vessels are intriguing and various analyses indicate their bidirectional signals in multiple aspects of cancer development and progression (69). The GO analysis revealed aspartic-type peptidase activity to be associated with LUAD and LUSC progressions, that is, Napsin A, expressed in the lung cells to process surfactant protein B (SP-B) which may be associated with *CKS1B* and the correlated genes (70). Moreover, the TCF/β-catenin complex can initiate proto-oncogenic transcriptions leading to activated transcription of downstream cell-proliferation-related genes. This process helps in cell proliferation and therefore, the abnormal β-catenin expression indicates malignancies (71). The pathways and GO enrichment analysis exhibited *CKS1B* significance and that of its correlated genes in varying oncogenic processes promoting LC progression (**Figure 10).** While these extensive evaluations suggest the use of *CKS1B* gene expression as prognostic biomarkers for the early diagnosis of human LUADs and LUSCs, further in vivo and in vitro studies are required to determine the final application approaches.

## 5. Conclusion

LC is the predominant cause of cancer-related deaths worldwide. This study aimed to identify the molecular signatures involved in the development and progression of LUAD and LUSC. Cancer prognostic factors are essential for early diagnosis and effective treatment, reducing the risk of overtreatment for patients. *CKS1B* appears to be involved in both LUAD and LUSC, according to our findings. The study further points to the possibility of using *CKS1B* as a biomarker for early LC detection and medication screening in treatment strategies. Furthermore, since *CKS1B* is involved in several important signaling pathways, inhibiting it may be a new way to prevent or reduce LC. *CKS1B* and its expression in LC progression were also linked to potential signaling mechanisms and gene ontological characteristics, according to the study. Researchers could be able to stop cancer from spreading by working on these pathways. As a result, *CKS1B* has been suggested as a useful biomarker and potential therapeutic target for human LC regulation or prevention.

## Acknowledgements

Authors are thankful to the Department of Genetic Engineering and Biotechnology, University of Chittagong, Chattogram, Bangladesh and International Foundation for Collaborative Research, Chattogram, Bangladesh, a dynamic platform of young researchers, for providing support to successfully carry out the research.

## 6. Declarations Ethics declarations

Not Applicable

## Approval for human/animal experiments

Not Applicable

## Consent for participation/publication

Not Applicable

## Availability of data and material

All the data generated during the experiment are provided in the manuscript/supplementary material.

## References

1. What Is Cancer? (2021). Retrieved 19 March 2021, from https://www.cancer.gov/about-cancer/understanding/what-is-cancer

2. Sung, H., Ferlay, J., Siegel, R., Laversanne, M., Soerjomataram, I., Jemal, A., & Bray, F. (2021). Global cancer statistics 2020: GLOBOCAN estimates of incidence and mortality worldwide for 36 cancers in 185 countries. CA: A Cancer Journal for Clinicians. DOI: https://doi.org/10.3322/caac.21660 PMid:33538338

3. What Is Lung Cancer? | Types of Lung Cancer. (2021). Retrieved 19 March 2021, from https://www.google.com/amp/s/amp.cancer.org/cancer/lung-cancer/about/what-is.html

4. Bender, E. (2014). Epidemiology: The dominant malignancy. Nature, 513(7517), S2–S3. DOI: https://doi.org/10.1038/513S2a PMid:25208070

5. Foss, K., Sima, C., Ugolini, D., Neri, M., Allen, K., & Weiss, G. (2011). miR-1254 and miR-574-5p: Serum-Based microRNA Biomarkers for Early-Stage Non-small Cell Lung Cancer. Journal of Thoracic Oncology, 6(3), 482–488. DOI: https://doi.org/10.1097/JTO.0b013e318208c785 PMid:21258252

6. Wong, M., Lao, X., Ho, K., Goggins, W., & Tse, S. (2017). Incidence and mortality of lung cancer: global trends and association with socioeconomic status. Scientific Reports, 7(1). DOI: https://doi.org/10.1038/s41598-017-14513-7 PMid:29085026

7. Jemal, A., Bray, F., Center, M., Ferlay, J., Ward, E., & Forman, D. (2011). Global cancer statistics. CA: A Cancer Journal for Clinicians, 61(2), 69–90. DOI: https://doi.org/10.3322/caac.20107 PMid:21296855

8. Yu, Y., & Tian, X. (2020). Analysis of genes associated with prognosis of lung adenocarcinoma based on GEO and TCGA databases. Medicine, 99(19). DOI: https://doi.org/10.1097/MD.0000000000020183 PMid:32384511

9. Shi, W., Huang, Q., Xie, J., Wang, H., Yu, X., & Zhou, Y. (2020). CKS1B as Drug Resistance-Inducing Gene-A Potential Target to Improve Cancer Therapy. Frontiers in Oncology, 10, 1978. DOI: https://doi.org/10.3389/fonc.2020.582451 PMid:33102238 PMCid:PMC7545642

10. Krishnan, A., Nair, S., & Pillai, M. (2009). Loss of cks1 homeostasis deregulates cell division cycle. Journal Of Cellular And Molecular Medicine, 14(1-2), 154–164. DOI: https://doi.org/10.1111/j.1582-4934.2009.00698.x PMid:19228269 PMCid:PMC3837597

11. Stella, F., Pedrazzini, E., Baialardo, E., Fantl, D., Schutz, N., & Slavutsky, I. (2014). Quantitative analysis of CKS1B mRNA expression and copy number gain in patients with plasma cell disorders. Blood Cells, Molecules, And Diseases, 53(3), 110–117. DOI: https://doi.org/10.1016/j.bcmd.2014.05.006 PMid:24973170

12. Zeng, Z., Gao, Z., Zhang, Z., Jiang, H., Yang, C., Yang, J., & Xia, X. (2019). Downregulation of CKS1B restrains the proliferation, migration, invasion and angiogenesis of retinoblastoma cells through the MEK/ERK signaling pathway. International Journal Of Molecular Medicine. DOI: https://doi.org/10.3892/ijmm.2019.4183

13. Lee, E., Kim, D., Kim, J., & Yoon, Y. (2011). Cell-Cycle Regulator Cks1 Promotes Hepatocellular Carcinoma by Supporting NF-κB-Dependent Expression of Interleukin-8. Cancer Research, 71(21), 6827–6835. DOI: https://doi.org/10.1158/0008-5472.CAN-10-4356 PMid:21917729

14. Zhan, F., Colla, S., Wu, X., Chen, B., Stewart, J., & Kuehl, W. et al. (2007). CKS1B, overexpressed in aggressive disease, regulates multiple myeloma growth and survival through SKP2- and p27Kip1-dependent and -independent mechanisms. Blood, 109(11), 4995–5001. DOI: https://doi.org/10.1182/blood-2006-07-038703

15. Wang, X., Xu, J., Ju, S., Ni, H., Zhu, J., & Wang, H. (2010). Livin gene plays a role in drug resistance of colon cancer cells. Clinical Biochemistry, 43(7-8), 655–660. DOI: 0.1016/j.clinbiochem.2010.02.004 PMid: 20171199

16. Zolota, V., Tzelepi, V., Leotsinidis, M., Zili, P., Panagopoulos, N., & Dougenis, D. et al. (2010). Histologic-Type Specific Role of Cell Cycle Regulators in Non-Small Cell Lung Carcinoma. Journal Of Surgical Research, 164(2), 256–265. DOI: https://doi.org/10.1016/j.jss.2009.03.035 PMid:19691991

17. Kitajima, S., Kudo, Y., Ogawa, I., Bashir, T., Kitagawa, M., & Miyauchi, M. et al. (2004). Role of Cks1 Overexpression in Oral Squamous Cell Carcinomas. The American Journal Of Pathology, 165(6), 2147–2155. DOI: https://doi.org/10.1016/S0002-9440(10)63264-6

18. Slotky, M., Shapira, M., Ben-Izhak, O., Linn, S., Futerman, B., Tsalic, M., & Hershko, D. (2005). The expression of the ubiquitin ligase subunit Cks1 in human breast cancer. Breast Cancer Research, 7(5). DOI: https://doi.org/10.1186/bcr1278 PMid:16168119 PMCid:PMC1242136

19. Shaughnessy, J., Zhan, F., Burington, B., Huang, Y., Colla, S., & Hanamura, I. et al. (2006). A validated gene expression model of high-risk multiple myeloma is defined by deregulated expression of genes mapping to chromosome 1. Blood, 109(6), 2276–2284. DOI: https://doi.org/10.1182/blood-2006-07-038430 PMid:17105813

20. Chen, M., Qi, C., Reece, D., & Chang, H. (2012). Cyclin kinase subunit 1B nuclear expression predicts an adverse outcome for patients with relapsed/refractory multiple myeloma treated with bortezomib. Human Pathology, 43(6), 858–864. DOI: https://doi.org/10.1016/j.humpath.2011.07.013 PMid:22047644

21. Huang, J., Zhou, Y., Thomas, G., Gu, Z., Yang, Y., & Xu, H. et al. (2015). NEDD8 Inhibition Overcomes CKS1B-Induced Drug Resistance by Upregulation of p21 in Multiple Myeloma. Clinical Cancer Research, 21(24), 5532–5542. DOI: https://doi.org/10.1158/1078-0432.CCR-15-0254 PMid:26156395 PMCid:PMC4804624

22. Hwang, J. S., Jeong, E. J., Choi, J., Lee, Y. J., Jung, E., Kim, S. K., … & Kim, J. S. (2019). MicroRNA-1258 inhibits the proliferation and migration of human colorectal cancer cells through suppressing CKS1B expression. Genes, 10(11), 912. DOI: https://doi.org/10.3390/genes10110912 PMid:31717435 PMCid:PMC6896137

23. Lemjabbar-Alaoui, H., Hassan, O., Yang, Y., & Buchanan, P. (2015). Lung cancer: Biology and treatment options. Biochimica Et Biophysica Acta (BBA) - Reviews On Cancer, 1856(2), 189–210. DOI: https://doi.org/10.1016/j.bbcan.2015.08.002 PMid:26297204 PMCid:PMC4663145

24. Tang Z, Kang B, Li C, Chen T, Zhang Z. GEPIA2: an enhanced web server for large-scale expression profiling and interactive analysis. Nucleic acids research. 2019 Jul 2;47(W1):W556–60. DOI: https://doi.org/10.1093/nar/gkz430 PMid:31114875 PMCid:PMC6602440

25. Park SJ, Yoon BH, Kim SK, Kim SY. GENT2: an updated gene expression database for normal and tumor tissues. BMC medical genomics. 2019 Jul;12(5):101. DOI: https://doi.org/10.1186/s12920-019-0514-7 PMid:31296229 PMCid:PMC6624177

26. Rhodes, D.R., Yu, J., Shanker, K., Deshpande, N., Varambally, R., Ghosh, D., Barrette, T., Pandey, A. and Chinnaiyan, A.M., 2004. ONCOMINE: a cancer microarray database and integrated data-mining platform. Neoplasia (New York, NY), 6(1), p.1. DOI: https://doi.org/10.1016/S1476-5586(04)80047-2

27. Chandrashekar DS, Bashel B, Balasubramanya SA, Creighton CJ, Ponce-Rodriguez I, Chakravarthi BV, Varambally S. UALCAN: a portal for facilitating tumor subgroup gene expression and survival analyses. Neoplasia. 2017 Aug 1;19(8):649–58. DOIDOI: https://doi.org/10.1016/j.neo.2017.05.002 PMid:28732212 PMCid:PMC5516091

28. Uhlén M, Fagerberg L, Hallström BM, Lindskog C, Oksvold P, Mardinoglu A, Sivertsson Å, Kampf C, Sjöstedt E, Asplund A, Olsson I. Tissue-based map of the human proteome. Science. 2015 Jan 23;347(6220). DOI: 10.1126/science.1260419 DOI: https://doi.org/10.1126/science.1260419 PMid:25613900

29. Goldman MJ, Craft B, Hastie M, Repečka K, McDade F, Kamath A, Banerjee A, Luo Y, Rogers D, Brooks AN, Zhu J. Visualizing and interpreting cancer genomics data via the Xena platform. Nature Biotechnology. 2020 May 22:1–4. DOI: https://doi.org/10.1038/s41587-020-0546-8 PMid:32444850 PMCid:PMC7386072

30. Cerami, E., Gao, J., Dogrusoz, U., Gross, B.E., Sumer, S.O., Aksoy, B.A., Jacobsen, A., Byrne, C.J., Heuer, M.L., Larsson, E. and Antipin, Y., 2012. The cBio cancer genomics portal: an open platform for exploring multidimensional cancer genomics data. DOI: https://doi.org/10.1158/2159-8290.CD-12-0095 PMid:22588877 PMCid:PMC3956037

31. Mizuno H, Kitada K, Nakai K, Sarai A. PrognoScan: a new database for meta-analysis of the prognostic value of genes. BMC medical genomics. 2009 Dec 1;2(1):18. DOI: https://doi.org/10.1186/1755-8794-2-18 PMid:19393097 PMCid:PMC2689870

32. Chen EY, Tan CM, Kou Y, Duan Q, Wang Z, Meirelles GV, Clark NR, Ma’ayan A. Enrichr: interactive and collaborative HTML5 gene list enrichment analysis tool. BMC bioinformatics. 2013 Dec 1;14(1):128. DOI: https://doi.org/10.1186/1471-2105-14-128 PMid:23586463 PMCid:PMC3637064

33. Kuleshov MV, Jones MR, Rouillard AD, Fernandez NF, Duan Q, Wang Z, Koplev S, Jenkins SL, Jagodnik KM, Lachmann A, McDermott MG. Enrichr: a comprehensive gene set enrichment analysis web server 2016 update. Nucleic acids research. 2016 Jul 8;44(W1):W90–7. DOI: https://doi.org/10.1093/nar/gkw377 PMid:27141961 PMCid:PMC4987924

34. Cancer Genome Atlas Research Network. Comprehensive molecular profiling of lung adenocarcinoma. Nature, 511 (2014), pp. 543-550 DOI: https://doi.org/10.1038/nature13385 PMid:25079552 PMCid:PMC4231481

35. Yang, Y., Wang, M., & Liu, B. (2019). Exploring and comparing of the gene expression and methylation differences between lung adenocarcinoma and squamous cell carcinoma. Journal of cellular physiology, 234(4), 4454–4459. DOI: https://doi.org/10.1002/jcp.27240 PMid:30317601

36. Cruz, C. S. D., Tanoue, L. T., & Matthay, R. A. (2011). Lung cancer: epidemiology, etiology, and prevention. Clinics in chest medicine, 32(4), 605–644. DOI: https://doi.org/10.1016/j.ccm.2011.09.001 PMid:22054876 PMCid:PMC3864624

37. Bernicker, E. H., Allen, T. C., & Cagle, P. T. (2019). Update on emerging biomarkers in lung cancer. Journal of thoracic disease, 11(Suppl 1), S81. DOI: https://doi.org/10.21037/jtd.2019.01.46 PMid:30775031 PMCid:PMC6353743

38. K.G.M. Moons, D.G. Altman, Y. Vergouwe, P. Royston, Prognosis and prognostic research: application and impact of prognostic models in clinical practice, BMJ (Clinical research ed.) 338 (2009) b606, b606. DOI: https://doi.org/10.1136/bmj.b606 PMid:19502216

39. Walters S, et al. 2013. Lung cancer survival and stage at diagnosis in Australia, Canada, Denmark, Norway, Sweden and the UK: a population-based study, 2004-2007. Thorax. 68, 551-564. DOI: https://doi.org/10.1136/thoraxjnl-2012-202297 PMid:23399908

40. Shah R, Sabanathan S, Richardson J, Mearns AJ, Goulden C. 1996. Results of surgical treatment of stage I and II lung cancer. J. Cardiovasc. Surg. 37, 169–172.

41. Nesbitt JC, Putnam JB Jr, Walsh GL, Roth JA, Mountain CF. 1995. Survival in early-stage non-small cell lung cancer. Ann. Thorac. Surg. 60, 466–472. DOI: https://doi.org/10.1016/0003-4975(95)00169-L

42. Goldstraw P, et al. 2016. The IASLC lung cancer staging project: proposals for revision of the TNM stage groupings in the forthcoming (eighth) edition of the TNM classification for lung cancer. J. Thorac. Oncol. 11, 39–51. DOI: https://doi:10.1016/j.jtho.2015.09.009

43. Bradley, S. H., Kennedy, M. P., & Neal, R. D. (2019). Recognising lung cancer in primary care. Advances in therapy, 36(1), 19–30. DOI: https://doi.org/10.1007/s12325-018-0843-5 PMid:30499068 PMCid:PMC6318240

44. Shi W, Huang Q, Xie J, Wang H, Yu X, Zhou Y. CKS1B as Drug Resistance-Inducing Gene-A Potential Target to Improve Cancer Therapy. Front Oncol. 2020 Sep 25;10:582451. doi: 10.3389/fonc.2020.582451. PMID: 33102238; PMCID: PMC7545642.

45. Laurinaviciene, V. Ostapenko, D. Dasevicius, S. Jarmalaite, J. Lazutka, Immunohistochemistry profiles of breast ductal carcinoma: factor analysis of digital image analysis data, Diagn. Pathol. 7 (2012) 27, 27. DOI: https://doi.org/10.1186/1746-1596-7-27 PMid:22424533 PMCid:PMC3319425

46. Laurinavicius, A., Plancoulaine, B., Herlin, P., & Laurinaviciene, A. (2016). Comprehensive immunohistochemistry: digital, analytical and integrated. Pathobiology, 83(2-3), 156–163. DOI: https://doi.org/10.1159/000442389 PMid:27101138

47. Yu Y, Tian X. Analysis of genes associated with prognosis of lung adenocarcinoma based on GEO and TCGA databases. Medicine (Baltimore). 2020 May;99(19):e20183. doi: 10.1097/MD.0000000000020183. PMID: 32384511; PMCID: PMC7220259.

48. Liu, L., Yang, Y., Liu, S., Tao, T., Cai, J., Wu, J., … & Li, M. (2019). EGF-induced nuclear localization of SHCBP1 activates β-catenin signaling and promotes cancer progression. Oncogene, 38(5), 747–764. DOI: https://doi.org/10.1038/s41388-018-0473-z PMid:30177836 PMCid:PMC6355651

49. Santamaria D, Barriere C, Cerqueira A, et al. Cdk1 is sufficient to drive the mammalian cell cycle. Nature 2007; 448: 811–815. DOI: https://doi.org/10.1038/nature06046 PMid:17700700

50. Malumbres M and Barbacid M. Cell cycle, CDKs and cancer: a changing paradigm. Nat Rev Cancer 2009; 9: 153-166. DOI: https://doi.org/10.1038/nrc2602 PMid:19238148

51. Enserink JM and Kolodner RD. An overview of Cdk1-controlled targets and processes. Cell Div 2010; 5: 11. DOI: https://doi.org/10.1186/1747-1028-5-11 PMid:20465793 PMCid:PMC2876151

52. Ravindran Menon D, Luo Y, Arcaroli JJ, et al. CDK1 interacts with Sox2 and promotes tumor initiation in human melanoma. Cancer Res 2018; 78: 6561–6574. DOI: https://doi.org/10.1158/0008-5472.CAN-18-0330 PMid:30297536 PMCid:PMC6279496

53. Li, M., He, F., Zhang, Z., Xiang, Z., & Hu, D. (2020). CDK1 serves as a potential prognostic biomarker and target for lung cancer. Journal of International Medical Research, 48(2), 0300060519897508. DOI: https://doi.org/10.1177/0300060519897508 PMid:32020821 PMCid:PMC7111107

54. You, H., Lin, H., & Zhang, Z. (2015). CKS2 in human cancers: Clinical roles and current perspectives. Molecular and clinical oncology, 3(3), 459–463. DOI: https://doi.org/10.3892/mco.2015.501 PMid:26137251 PMCid:PMC4471627

55. Weirauch, M. T. (2011). Gene coexpression networks for the analysis of DNA microarray data. Applied statistics for network biology: methods in systems biology, 1, 215–250. DOI: https://doi.org/10.1002/9783527638079.ch11

56. Melak, T., & Gakkhar, S. (2015). Comparative genome and network centrality analysis to identify drug targets of mycobacterium tuberculosis H37Rv. BioMed research international, 2015. DOI: https://doi.org/10.1155/2015/212061 PMid:26618166 PMCid:PMC4651637

57. Gursoy, A., Keskin, O., & Nussinov, R. (2008). Topological properties of protein interaction networks from a structural perspective. Biochemical Society Transactions, 36(6), 1398–1403. DOI: https://doi.org/10.1042/BST0361398 PMid:19021563 PMCid:PMC7243876

58. McCubrey, J. A., Steelman, L. S., Chappell, W. H., Abrams, S. L., Wong, E. W., Chang, F., … & Franklin, R. A. (2007). Roles of the Raf/MEK/ERK pathway in cell growth, malignant transformation and drug resistance. Biochimica et Biophysica Acta (BBA)-Molecular Cell Research, 1773(8), 1263–1284. DOI: https://doi.org/10.1016/j.bbamcr.2006.10.001 PMid:17126425 PMCid:PMC2696318

59. Desai, T. J., Brownfield, D. G., & Krasnow, M. A. (2014). Alveolar progenitor and stem cells in lung development, renewal and cancer. Nature, 507(7491), 190–194. DOI: https://doi.org/10.1038/nature12930 PMid:24499815 PMCid:PMC4013278

60. Herbst, R. S., Morgensztern, D., & Boshoff, C. (2018). The biology and management of non-small cell lung cancer. Nature, 553(7689), 446–454. DOI: https://doi.org/10.1038/nature25183 PMid:29364287

61. J.D. Campbell, A. Alexandrov, J. Kim, J. Wala, A.H. Berger, C.S. Pedamallu, S.A. Shukla, G. Guo, A.N. Brooks, B.A. Murray, et al., Cancer Genome Atlas Research Network Distinct patterns of somatic genome alterations in lung adenocarcinomas and squamous cell carcinomas. Nat. Genet., 48 (2016), pp. 607–616 DOI: https://doi.org/10.1038/ng.3564 PMid:27158780 PMCid:PMC4884143

62. Desai, A., Menon, S. P., & Dy, G. K. (2016). Alterations in genes other than EGFR/ALK/ROS1 in non-small cell lung cancer: trials and treatment options. Cancer biology & medicine, 13(1), 77. DOI: https://doi.org/10.20892/j.issn.2095-3941.2016.0008

63. Beigel, J. H., Tomashek, K. M., Dodd, L. E., Mehta, A. K., Zingman, B. S., Kalil, A. C., … & Lane, H. C. (2020). Remdesivir for the treatment of Covid-19-preliminary report. The New England journal of medicine. DOI: https://doi.org/10.1056/NEJMoa2007764

64. Saini, R., Batra, U., Jain, A., & Agrawal, C. (2016). EGFR mutation--a commonly neglected mutation in squamous cell lung carcinoma. Journal of Cancer Metastasis and Treatment, 2, 253–254. DOI: https://doi.org/10.20517/2394-4722.2016.07

65. Bethune, G., Bethune, D., Ridgway, N., & Xu, Z. (2010). Epidermal growth factor receptor (EGFR) in lung cancer: an overview and update. Journal of thoracic disease, 2(1), 48.

66. Tan, A. C. (2020). Targeting the PI3K/AKT/mTOR pathway in non small cell lung cancer (NSCLC). Thoracic cancer, 11(3), 511–518. DOI: https://doi.org/10.1111/1759-7714.13328 PMid:31989769 PMCid:PMC7049515

67. Yuan, T. L., and Cantley, L. C. (2008). PI3K pathway alterations in cancer: variations on a theme. Oncogene 27, 5497–5510. doi: 10.1038/onc.2008.245 DOI: https://doi.org/10.1038/onc.2008.245 PMid:18794884 PMCid:PMC3398461

68. Scheffler M, Bos M, Gardizi M et al PIK3CA mutations in non small cell lung cancer (NSCLC): Genetic heterogeneity, prognostic impact and incidence of prior malignancies. Oncotarget 2015; 6: 1315–26. DOI: https://doi.org/10.18632/oncotarget.2834 PMid:25473901 PMCid:PMC4359235

69. Pasquale, E. B. (2010). Eph receptors and ephrins in cancer: bidirectional signalling and beyond. Nature Reviews Cancer, 10(3), 165–180. DOI: https://doi.org/10.1038/nrc2806 PMid:20179713 PMCid:PMC2921274

70. Ueno, T., Elmberger, G., Weaver, T. E., Toi, M., & Linder, S. (2008). The aspartic protease napsin A suppresses tumor growth independent of its catalytic activity. Laboratory investigation, 88(3), 256–263. DOI: https://doi.org/10.1038/labinvest.3700718 PMid:18195689

71. Gao, C., Wang, Y., Broaddus, R., Sun, L., Xue, F., & Zhang, W. (2018). Exon 3 mutations of CTNNB1 drive tumorigenesis: a review. Oncotarget, 9(4), 5492. DOI: https://doi.org/10.18632/oncotarget.23695 PMid:29435196 PMCid:PMC5797067

